# Engineering Rotating Apical-Out Airway Organoid for Assessing Respiratory Cilia Motility

**DOI:** 10.1101/2022.01.15.476455

**Authors:** Piyumi Wijesekara, Prakarsh Yadav, Lydia A. Perkins, Donna B. Stolz, Jonathan M. Franks, Simon C. Watkins, Emily Reinoso Jacome, Steven L. Brody, Amjad Horani, Jian Xu, Amir Barati Farimani, Xi Ren

**Affiliations:** Department of Biomedical Engineering, Carnegie Mellon University, 5000 Forbes Avenue, Pittsburgh, Pennsylvania, United States; Department of Mechanical Engineering, Carnegie Mellon University, 5000 Forbes Avenue, Pittsburgh, Pennsylvania, United States; Department of Biological Sciences, Carnegie Mellon University, 5000 Forbes Avenue, Pittsburgh, Pennsylvania, United States; Department of Cell Biology, University of Pittsburgh School of Medicine, Pittsburgh, Pennsylvania, United States; Department of Medicine, Washington University School of Medicine, St. Louis, Missouri, United States; Department of Pediatrics, Washington University School of Medicine, St. Louis, Missouri, United States; Department of Cell Biology and Physiology, Washington University School of Medicine, St. Louis, Missouri, United States

## Abstract

Motile cilia project from the airway apical surface and directly interface with inhaled external environment. Due to cilia’s nanoscale dimension and high beating frequency, quantitative assessment of their motility remains a sophisticated task. Here we described a robust approach for reproducible engineering of apical-out airway organoid (AOAO) of defined size. Propelled by exterior-facing cilia beating, the mature AOAO exhibited stable rotational motion when surrounded by Matrigel. We developed a computational framework leveraging computer vision algorithms to quantify AOAO rotation and validated its correlation with direct measurement of cilia motility. We further established the feasibility of using AOAO rotation to recapitulate and measure defective cilia motility caused by chemotherapy-induced toxicity and by CCDC39 mutations in cells from primary ciliary dyskinesia patient. We expect our rotating AOAO model and the associated computational pipeline to offer a generalizable framework to expediate modeling of and therapeutic development for genetic and environmental ciliopathies.

## Introduction

Motile cilia are specialized, highly conserved organelles that project from the luminal epithelial surface lining the respiratory tract, middle ear cavity, fallopian tube, and brain ventricles.^1-8^ Motile cilia have the typical 9+2 microtubule architecture with a central pair of microtubule singlets surrounded by nine outer microtubule doublets.^9,10^ Motile cilia function as mechanical nanomachines that generate high-speed beating motion from cycles of unidirectional sliding of outer doublet microtubules, which are powered by dynein molecular motors.^9,11,12^ Coordinated cilia beating serves critical functions in facilitating the directional transport of luminal substances, such as mucus in the respiratory tract and fertilized egg in the fallopian tube.^1,2,5,6^

Abnormal cilia motility (ciliopathies) can result from genetic disorders affecting the structure or function of motile cilia, such as primary ciliary dyskinesia (PCD), which lead to devastating consequences, including chronic infection in the lung and ear, laterality defects, infertility, and rarely abnormal accumulation of cerebrospinal fluid in the brain.^13-15^ There are no therapeutic cures that can reverse the defects in cilia motility or halt the progression of diseases caused by genetic cilia abnormalities in PCD.^13-15^ Acquired motile ciliopathies may result from inhaled or ingested ciliotoxic chemicals that have been extensively implicated to compromise cilia motility in the middle ear,^16,17^ fallopian tube,^18,19^ and brain ventricles.^20,21^ In the respiratory system alone, cilia dysfunction is a pathological finding observed in several chronic diseases, and especially cigarette smoke related, which together affect over 35 million Americans.^2,22^ Thus, understanding the fundamental mechanisms regulating cilia motility under various pathophysiological conditions is of pivotal importance.

Airway organoids engineered from patient stem cells are a promising model for investigating respiratory pathophysiology, including ciliopathies. The airway epithelium is polarized with cilia beating and mucus secretion taking place on its apical surface that directly interacts with the inhaled air. However, the nanoscale dimension (∼200 nm diameter) of cilia combined with their high beating frequency (10-40 Hz) makes the measurement of cilia motility and function a challenging task. Over the past decades, high-speed video microscopy has emerged as a powerful tool for quantitative imaging of individual cilium motion in live cells and tissues. However, it requires highly specialized microscopic setup and cumbersome analytical process,^23,24^ and is difficult to scale up to a high-throughput format. Thus, biomedical advancements are needed to enable comprehensive assessment and investigation of cilia pathophysiology and to promote effective therapeutic development.

Here we describe a suspension, hydrogel-free culture strategy for reproducible engineering of apical-out airway organoid of defined size. Importantly, powered by cilia beating on its exterior surface, the apical-out organoid rotated in soft supporting material, which inspired the use of organoid rotation as a functional readout of respiratory cilia motility. We developed a computational framework that leveraged computer vision algorithms to reliably calculate the angular velocity of apical-out organoid rotation and correlated it with direct measurement of cilia motility. To assess such correlation, we analyzed organoids treated with known chemical modulators of cilia motility as well as those engineered from bronchial epithelial cells derived from healthy and PCD patients.

## Results

### Engineering apical-out airway organoid

The interaction between epithelial cells and their surrounding extracellular matrix (ECM) plays instrumental roles in determining tissue polarity. Apical-in organoids are typically produced from airway epithelial cells in ECM-embedded culture, leading to recognition of the organoid’s exterior surface that faces the ECM to be basal-lateral and its interior surface to be apical.^25-29^ Here we assessed whether removal of ECM support during airway organoid biogenesis from defined number of human airway basal stem cells (hABSCs) can reverse the apical-basal recognition and epithelial polarity (Figure 1A). Bronchus-derived hABSCs were expanded in two-dimensional (2D) culture using expansion medium formulated based on Bronchial Epithelial Cell Growth Medium.^30^ To enable airway organoid formation, defined number (500) of hABSCs, dissociated from 3D expansion, were allowed to aggregate together on top of a cell-repellent surface in 96-well plate with no ECM support. Following overnight suspension culture in differentiation medium (PneumaCult-ALI), we observed effective spheroid formation followed by differentiation into ciliated airway organoid with apical-out polarity (cilia beating on the outer surface) by the end of week 3 (Supplementary Video 1). This was referred to as Apical-Out Airway Organoid (AOAO). Compared to using expansion medium for hABSC cell aggregation followed by transitioning to differentiation medium, the use of differentiation medium for both initial cell aggregation and subsequent differentiation was essential for maintaining spheroid tissue integrity (Supplementary Figure 1).

**Figure 1.**
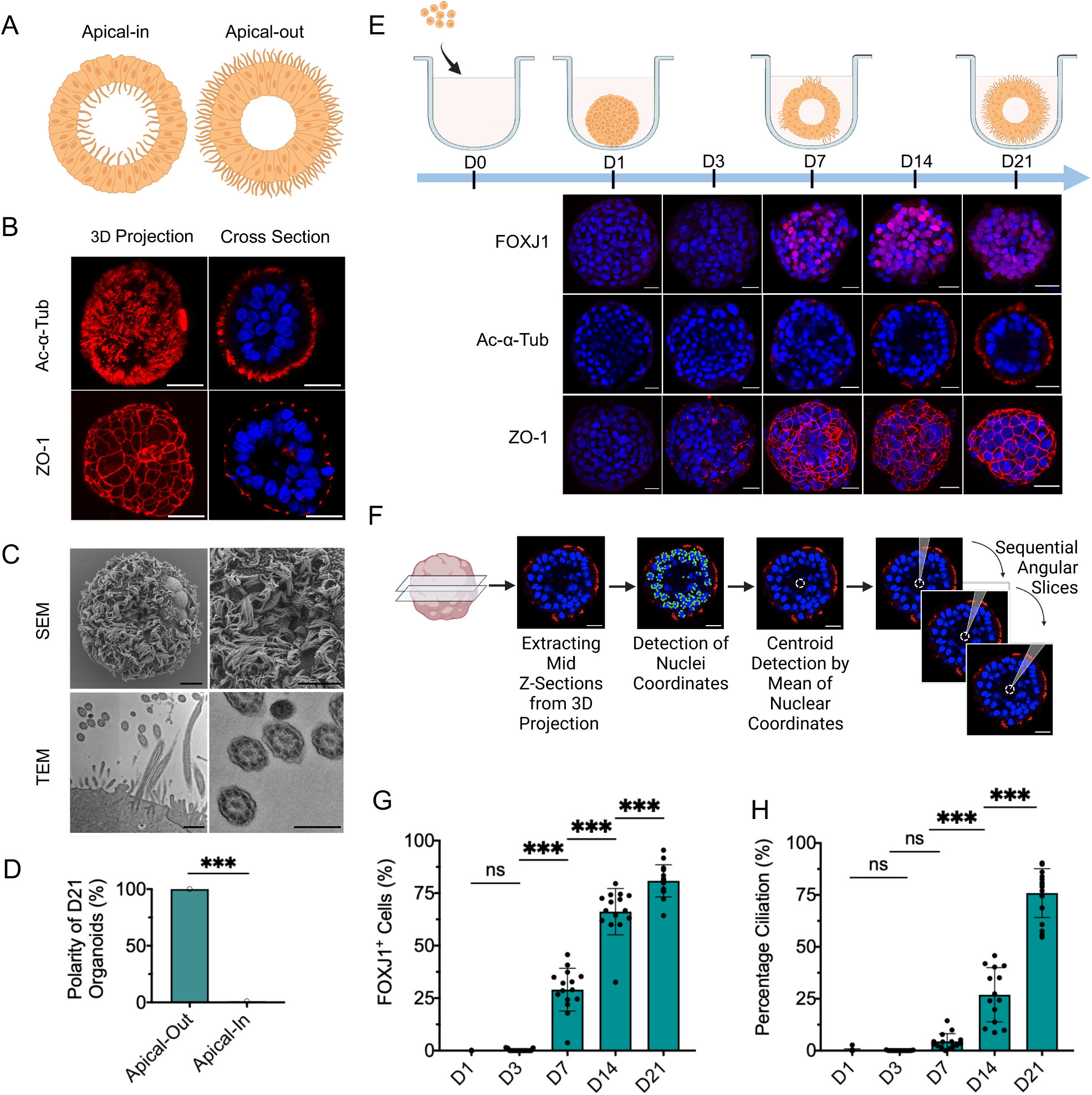
Characterization of the engineered AOAOs. (A) Diagram of apical-in versus apical-out airway organoids. (B) Immunofluorescence images of day-21 AOAOs stained with markers of cilia (Ac-α-Tub) and tight junction (ZO-1). Scale bar, 25 µm. (C) SEM (scale bar, 10 µm) and TEM (scale bar, 800 nm (left), 400 nm (right)) images of AOAOs. (D) Quantification of the percentage of day-21 (D21) organoids with apical-out versus apical-in epithelial polarity indicated by apical Ac-α-Tub localization. (E) Time-series characterization of AOAO maturation by immunostaining of FOXJ1 (nuclear marker of ciliated cells), Ac-α-Tub, and ZO-1. Scale bar, 25 µm. (F) Diagram showing the approach for assessing percentage ciliation by quantifying surface coverage of Ac-α-Tub expression. (G,H) Time-series quantification of FOXJ1^+^ cell abundance (G) and percentage ciliation (H) in AOAOs. All data represent means ± s.d. from ≥ 15 organoids. ***, *p*<0.001.

To characterize epithelial polarity in the resulting day-21 AOAOs, we performed immunofluorescence staining of key polarity markers of airway epithelium and observed highly selective localization of ciliary Acetylated-alpha-Tubulin (Ac-α-Tub) on the organoid outer surface. Consistent with this orientation, epithelial tight junction protein, Zona Occludens Protein-1 (ZO-1), formed highly organized intercellular junctions underneath the apical surface (Figure 1B). Using scanning electron microscopy (SEM) and transmission electron microscopy (TEM), we observed dense motile cilia covering the organoid outer surface with typical 9+2 microtubule organization, further verifying the apical-out epithelial polarity (Figure 1C). Next, we assessed the consistency of epithelial polarity in day-21 organoids resulting from continuous 3D suspension culture by examining Ac-α-Tub localization on the organoid’s exterior versus interior surface and observed 100% apical-out polarity (Figure 1D).

To track temporal dynamics of ciliogenesis and epithelial polarization, AOAOs were harvested on day-1, -3, -7, -14, and -21 of suspension differentiation, and evaluated for ciliated cell nuclear marker Forkhead Box J1 (FOXJ1), Ac-α-Tub, and ZO-1 (Figure 1E). FOXJ1^+^ ciliated cells emerged as early as day-7 and their abundance gradually increased to 81±8% on day-21 (Figure 1G). We then calculated the percentage ciliation by quantifying cilia coverage on the organoid’s exterior surface. To do this, mid-Z-sections were selected from confocal Z-stack images of each organoid whole-mount stained for Ac-α-Tub and Hoechst-33342 (nuclei). The centroid of each Z-section was identified using k-means clustering on nuclei coordinates. Angular slices of 1-degree from the centroid were assessed regarding their overlapping with ciliary Ac-α-Tub expression on the organoid edge. Finally, the percentage ciliation was calculated by normalizing the number of angular slices containing Ac-α-Tub fluorescence signal by 360 (Figure 1F). Applying this analytical pipeline to time-series images of AOAOs, we observed a steady increase in percentage ciliation over time, reaching 76±12% on day-21 (Figure 1H), which echoed the gradual increase in FOXJ1^+^ ciliated cell abundance (Figure 1G).

The native human airway is known to undergo goblet cell hyperplasia and mucus hypersecretion following stimulation with cytokines, such as Interleukin 13 (IL-13).^31^ In AOAOs engineered using standard differentiation medium, no MUC5AC^+^ goblet cells was observed on day-21. In sharp contrast, when IL-13 (5 ng/mL) was supplemented to the differentiation medium, massive induction of goblet cells was observed in day-21 AOAOs (Supplementary Figure 3).

### Assessing reversibility of AOAO epithelial polarity

Upon demonstrating the ECM-free, suspension culture as a driver for establishing consistent apical-out airway polarity, we next investigated the stability of such epithelial polarity when the surrounding extracellular environment changed. To do this, we transitioned hABSC aggregates, following only 1-day suspension culture, into Matrigel-embedded culture and continued the differentiation until day-21. To our surprise, as indicated by FOXJ1, Ac-α-Tub, and ZO-1 expression (Figure 2A), all organoids subjected to this two-phase culture procedure (1 day in suspension followed by 20 days in Matrigel embedding) remained exhibiting homogenous apical-out polarity (Figure 2B,C). Furthermore, these organoids from two-phase culture underwent robust ciliogenesis leading to day-21 ciliated cell abundance (FOXJ1^+^, 83±7%, Figure 2D) and percentage ciliation (70±14%, Figure 2E) that was not statistically different from that of standard AOAOs that have only experienced suspension culture (*p*=0.2964 for ciliated cell abundance; *p*=0.1448 for percentage ciliation). These results imply that airway epithelial polarity was effectively established within the first day of 3D suspension culture and remained stable even after being transitioned to ECM-supported culture. During the transition from suspension to Matrigel-embedded culture, we also observed sporadic merging of individual hABSC aggregates into larger organoid bodies, where Ac-α-Tub expression can be found on both the interior and exterior surfaces (Supplementary Figure 2).

**Figure 2.**
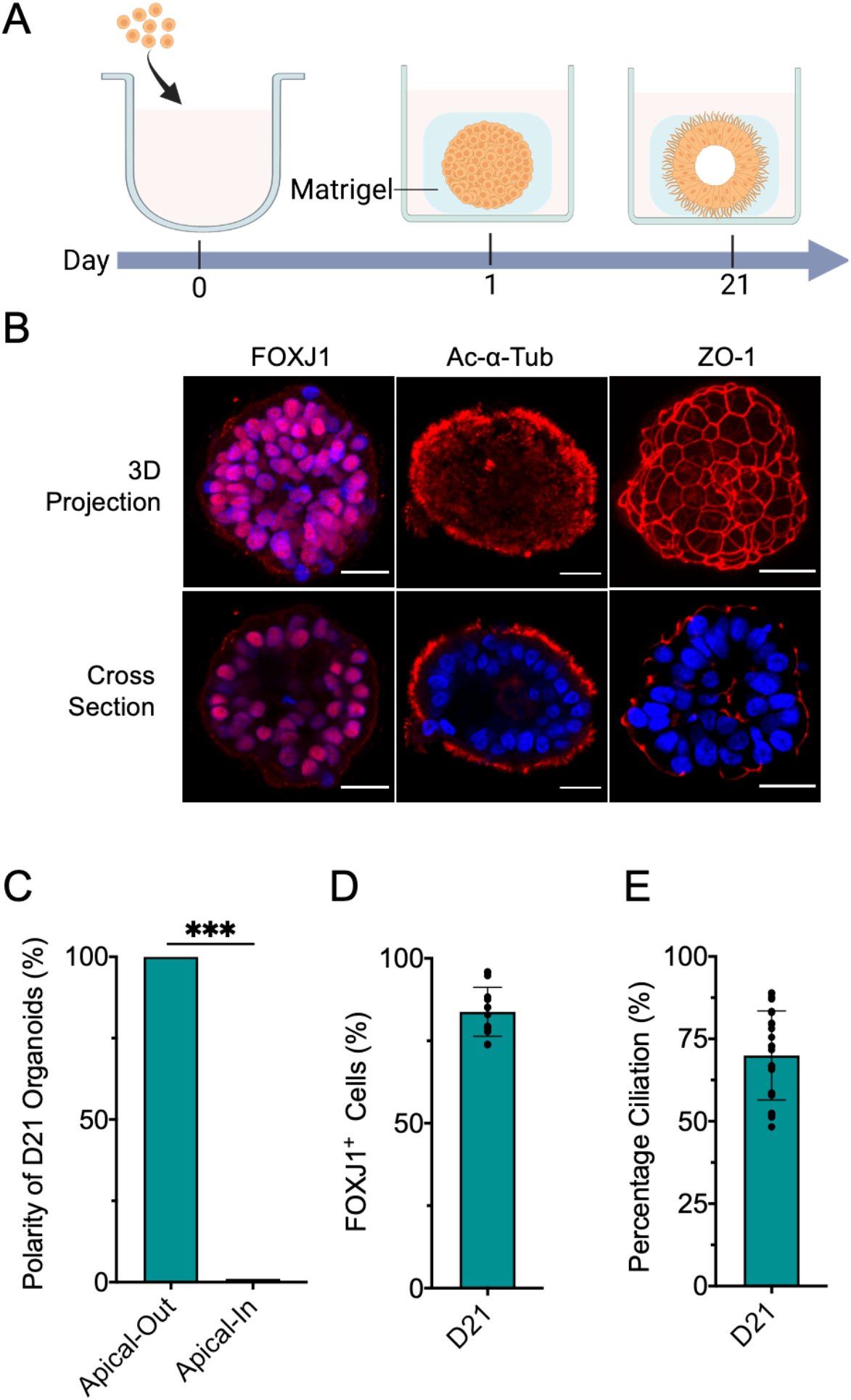
Assessing epithelial polarity reversibility in organoids transitioned from suspension to Matrigel-embedded culture. (A) Human ABSC aggregates were formed in ECM-free suspension culture for 1 day and then transferred into ECM-rich, Matrigel-embedded culture and maintained for an additional 20 days. (B) Immunofluorescence images of day-21 organoids stained for FOXJ1, Ac-α-Tub, and ZO-1. Scale bar, 25 µm. (C) Quantification of the percentage of day-21 (D21) organoids with apical-out versus apical-in epithelial polarity indicated by apical Ac-α-Tub localization. (D,E) Quantification of FOXJ1^+^ cell abundance (D) and percentage ciliation (E) in day-21 (D21) organoids. Data represent means ± s.d. from ≥ 15 organoids. ***, *p*<0.001.

### Developing computer vision algorithms to assess AOAO rotation

Intriguingly, beating motion of exterior-facing cilia endows motility to the AOAO, which exhibited random movement in suspension culture (Supplementary video 1). Here we investigated the possibility of stabilizing such cilia-powered AOAO motility by providing a 3D surrounding material support for cilia to propel against. To do this, mature AOAOs (between day-21 and day-28 of suspension differentiation) were embedded within Matrigel matrix (Figure 3A), which effectively enabled the AOAOs to adopt stable rotational motion (Supplementary Video 2), offering a unique opportunity to transform nanoscale, high-frequency cilia motility into microscale, low-frequency organoid rotation.

**Figure 3.**
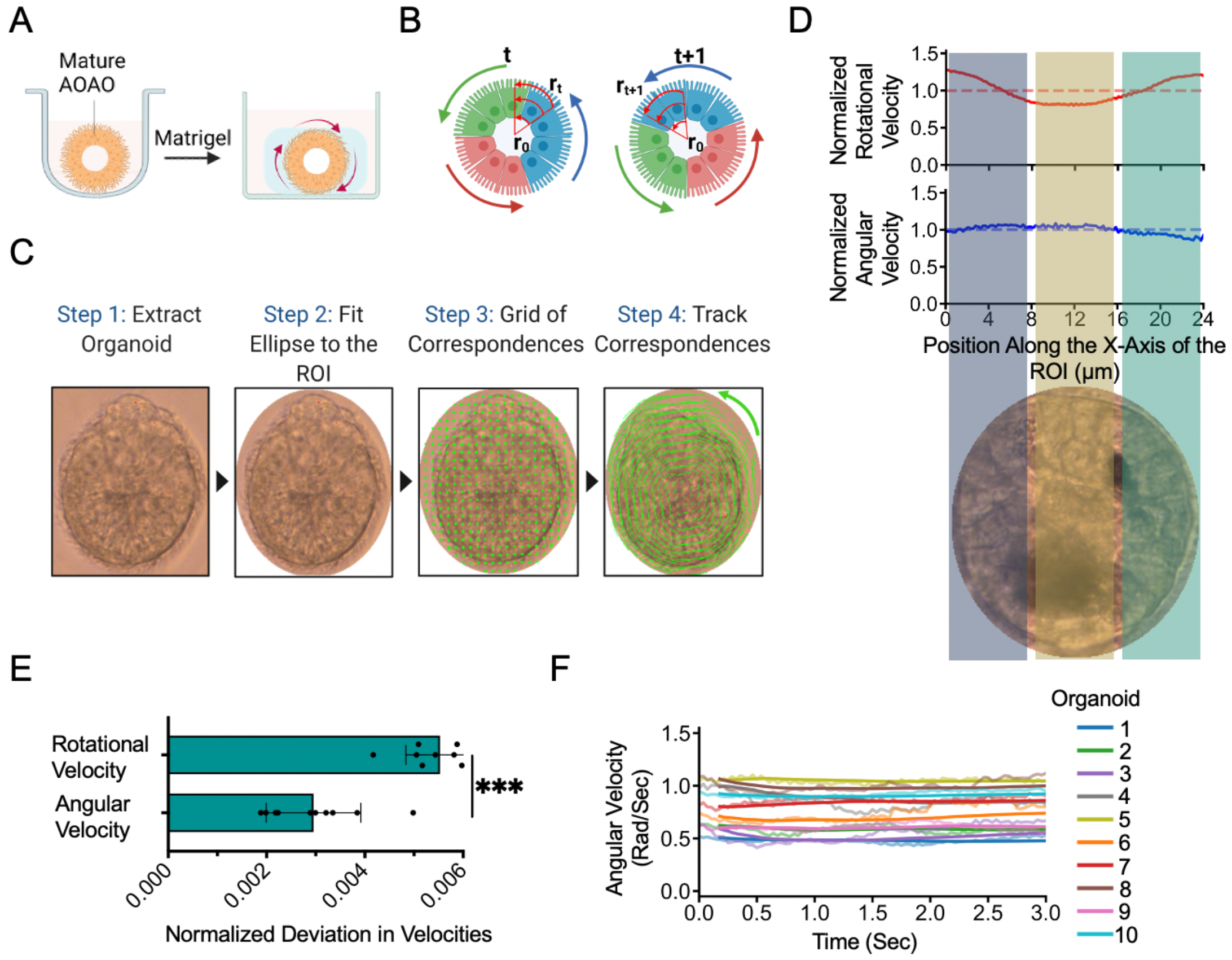
Enabling and quantifying AOAO rotation. (A) Diagram for enabling consistent AOAO rotational motion via matrix embedding. (B) Diagram depicting the computational method for calculating the organoid’s angular rotational motion: r0, the center of the organoid; rt, the position of the correspondence at time t; and rt+1, the position of the correspondence at time t+1. (C) Stepwise description of the computational framework used to calculate correspondence movement. (D) The rotational velocity (µm/sec) and angular velocity (rad/sec) normalized by their mean values for a representative organoid. The X-axis is the position of correspondences along the X-axis of the ROI. The velocity of correspondences on the organoid, perpendicular to every position on the X-axis were projected onto the X-axis and averaged to obtain the rotational or angular velocity for that position. The velocities were then normalized by their mean values. The color shaded regions represent the region of organoid which was used to calculate the rotational and angular velocity. (E) Deviation in the angular and rotational velocity with respect to their mean values of 10 representative organoids from three independent replicates. The deviation was calculated by taking the mean squared difference between the velocity profile and its mean value. The deviation was then normalized by the mean value. (F) The instantaneous angular velocity profile of 10 representative organoids from three independent replicates. The running mean (window = 5) of instantaneous angular velocity was shown as the solid line. ***, *p*<0.001.

To reliably quantify the rotational motion of AOAOs, we developed computer vision-based motion tracking algorithms.^32,33^ From video recordings of AOAO rotation (Supplementary Video 2), we detected the center of each organoid (r_0_) and the position of the correspondence being tracked (r_t_) and used these vectors to determine the distance of the correspondence from the center. The change in position of correspondence (r_t+1_) was used in the next step to calculate the distance covered by the correspondence (Figure 3B). To quantify the rotational motion, we identified the region of interest (ROI) by fitting an ellipse to the organoid to suppress the surrounding background. We generated a grid of correspondences in the ROI which were then tracked by the tracking algorithm. The distance covered by correspondences was then divided by the time taken to determine rotational velocity (Figure 3C).

The rotational velocity calculated above was dependent on the distance of the correspondence being tracked from the AOAO center. This led to large variation in measurements obtained at different regions of the same organoid. For example, the organoid’s rotational velocity profile had a parabolic shape with minimum at the central region and maximum at the periphery (Figure 3D, Supplementary Figure 4A). This was due to the correspondences close to the organoid center not covering a large distance in comparison to those at the periphery, which as a result yielded a lower rotational velocity. To overcome this constraint, we further calculated the angular velocity of each correspondence, which became independent on its exact position within the organoid, by dividing the rotational velocity by the distance of each correspondence from the organoid center (Supplementary Figure 4B). The angular velocity of the entire AOAO was determined by taking the mean of the angular velocity of all the correspondences being tracked (Supplementary Video 3). To compare the rotational and angular velocity profiles across the entire length of the AOAO, we calculated the mean squared deviation of the velocity from its mean value and then normalized it by the mean value (Figure 3E). The deviation in rotational velocity was 2-fold greater than that in angular velocity. Therefore, to ensure consistency in measuring AOAO rotational motion, we utilized the angular velocity as the main readout. Finally, to detect the time-dependent variability in tracking AOAO rotation, we visualized the instantaneous angular velocity of 10 representative AOAOs. The running mean of instantaneous angular velocity showed consistent rotational motion for AOAO throughout the entire recorded time period (Figure 3F).

### Assessing drug-induced inhibition of cilia motility and AOAO rotation

To assess the correlation between cilia motility and cilia-powered AOAO rotational motion, we applied known chemical inhibitors of cilia motility to Matrigel-embedded, mature AOAOs (Figure 4A). EHNA (erythro-9-(2-hydroxy-3-nonyl)adenine) is an inhibitor of dynein, the molecular motor that powers axonemal doublet microtubule sliding and thus cilia beating.^34,35^ We used the computer vision-based motion tracking algorithms developed above to compute the angular velocity of the same organoid before and after EHNA treatment (Figure 4B, Supplementary Video 4). We introduced EHNA at a range of concentrations (0, 0.1, 0.3 and 1 mM) to mature AOAOs for 2 hours, and observed an EHNA-dose-dependent reduction in organoid angular velocity (Figure 4C). In parallel, we confirmed the inhibitory effect of EHNA on cilia beating frequency (CBF) using kymography analysis (Figure 4D).

**Figure 4.**
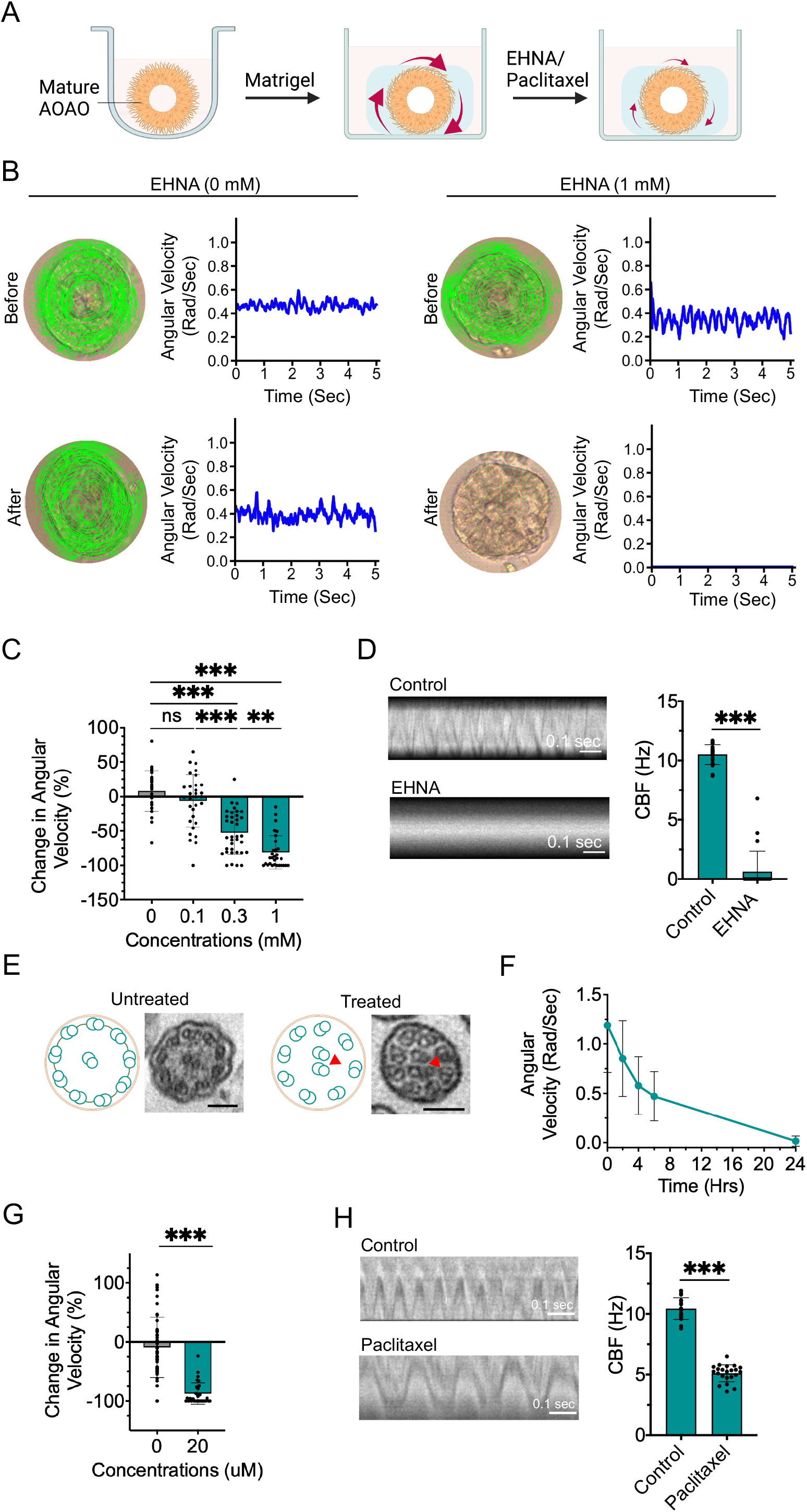
Characterization of AOAO rotation and cilia motility in the presence of cilia beating inhibitors. (A) Diagram showing AOAO rotational motion in response to EHNA or paclitaxel treatment. (B) Computational methods used for calculating organoid angular velocity following 1 mM EHNA treatment for 2 hours. (C) Quantification of organoid angular velocity following 2-hour treatment of EHNA at various concentrations. (D) CBF analyzed via conventional kymographs following 1 mM EHNA treatment for 2 hours. (E) TEM imaging of ciliary ultrastructure with and without 24-hour paclitaxel (20 µM) treatment. Scale bar, 100 nm. Arrowheads indicated mislocated microtubules. (F) Time-series analysis of paclitaxel’s effect on AOAO angular velocity. (G) Change in organoid angular velocity in the presence or absence of 20 µM paclitaxel treatment for 24 hours. (H) CBF quantification via kymograph analysis in the presence or absence of 20 µM Paclitaxel for 24 hours. Data represent means ± s.d. from three independent replicates with ≥ 10 organoids. **, *p*<0.01. ***, *p*<0.001.

Paclitaxel is a chemotherapeutic agent that stabilizes microtubule structures and thus interferes with microtubule-dependent mitosis, cell migration and cilia beating.^36-38^ Treatment of mature AOAOs with paclitaxel (20 µM) for 24 hours led to abnormalities in ciliary ultrastructure (Figure 4E), which is in line with prior studies.^39,40^ We then incubated the Matrigel-embedded AOAOs with paclitaxel (20 µM), monitored them periodically for 24 hours, and observed paclitaxel-induced, progressive reduction of organoid angular velocity (Figure 4F,G). Consistent with this, 24-hour paclitaxel treatment dramatically reduced CBF shown by kymography analysis (Figure 4H). These findings validated that the angular velocity of the AOAO correlates with and predicts cilia motility.

### Modeling and characterization of genetic ciliopathy using AOAOs

Primary ciliary dyskinesia (PCD) is a collection of genetic disorders involving abnormal motile cilia ultrastructure and function.^41-45^ Mutations in *CCDC39* gene cause inner dynein arm defects and axonemal disorganization in cilia and have been associated with PCD.^42^ Using hABSCs carrying mutations in *CCDC39* gene, we intended to assess whether AOAOs can be effectively generated from PCD-bearing cells and whether the PCD-associated ciliary defects can be recapitulated by the AOAO rotational motion. We expanded hABSCs isolated from healthy and PCD (with *CCDC39* mutations) patients and transitioned them for AOAO formation via 3D suspension culture (Figure 5A). Following 21 days of differentiation in suspension, as indicated by Ac-α-Tub and ZO-1 expression, airway organoids engineered from both healthy and PCD cells underwent effective epithelial differentiation with consistent apical-out polarity (Figure 5B). Further, we observed comparable percentage ciliation on the apical surface of healthy and PCD organoids (*p*=0.3212), indicating that the *CCDC39* mutations did not affect basic ciliogenesis (Figure 5C). However, as expected, PCD organoids exhibited defects in ciliary ultrastructure as indicated by TEM, showing a surrounding microtubule pair being mislocated to the center, compared to the normal 9+2 ciliary ultrastructure observed in healthy organoids (Figure 5D). Building on these morphological findings, we went on to assess and compare the rotational motion of PCD and healthy AOAOs by transferring them, following maturation, from 3D suspension culture to Matrigel embedding (Figure 5E). Consistent with defective ciliary structures, none of the embedded PCD AOAOs were able to rotate, as compared to over 75% of the embedded healthy AOAOs showing stable rotational motion (Figure 5F,G, and Supplementary Video 5). Lastly, we performed CBF analysis and did not observe obvious cilia motility in PCD organoids as compared to robust cilia beating in healthy organoids (Figure 5H). These findings further validated our AOAO model and its associated computational analysis pipeline as effective tools for modeling and assessing human motile ciliopathy.

**Figure 5.**
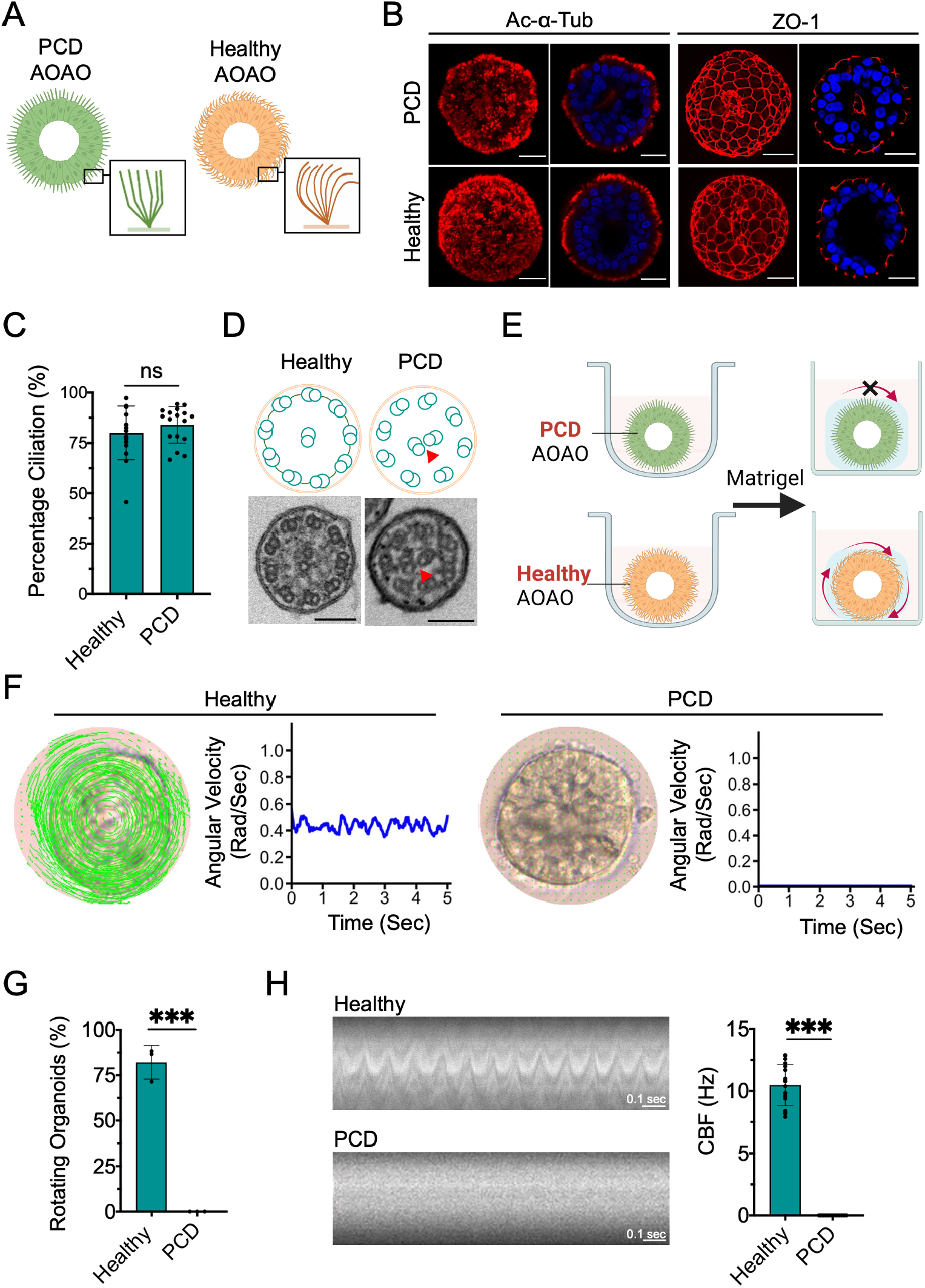
Modeling PCD-associated ciliary defects using AOAO rotation. (A) Diagram showing organoids engineered from hABSCs derived from PCD and healthy patients. (B) Imaging of day-21 AOAOs (PCD and healthy) stained for Ac-α-Tub and ZO-1. Scale bar, 25 µm. (C) Quantification of percentage ciliation in PCD and healthy AOAOs. (D) TEM images of healthy and PCD AOAOs showing cilia ultrastructural defects in the PCD organoids (arrowheads). Scale bar, 100 nm. (E) Diagram showing PCD and healthy AOAO’s rotational motion. (F) Computational methods used for calculating angular velocities of healthy and PCD AOAOs. (G) Rotation analysis of Matrigel-embedded, healthy and PCD AOAOs. (H) Kymograph-based CBF quantification of healthy and PCD AOAOs. ***, *p*<0.001.

## Discussion

In this study, we developed a robust approach to engineer apical-out airway organoids (AOAOs) with cilia beating taking placing at the organoid’s exterior surface, which effectively propelled the organoid to rotate when placed in 3D Matrigel support. We have established robust rotational velocity calculation algorithms and thereby a direct correlation between cilia motility and AOAO rotational motion. Our experimental results and computational tools will facilitate respiratory injury assessment with improved efficacy and will support the development of high-throughput assays for personalized disease management and therapeutic screening, benefiting patients suffering from respiratory diseases resulting from defective cilia function. Given the high-level similarity between motile cilia in different organ systems, we also expect wide applicability of our apical-out organoid model and its associated computational tools beyond the respiratory system.

In the native human lung, the apical airway surface is exposed directly to the external environment and therefore is the main interface interacting with respiratory pathogens, such as bacteria and viruses. The apical-out conformation enabled by the AOAO engineering described here will allow introduction of respiratory pathogens and pollutants directly to the apical airway surface in a non-invasive, repetitive manner with unprecedented convenience. While the apical-out epithelial polarity has previously been obtained via culturing epithelial sheets directly obtained from nasal biopsy in suspension culture, this method results in organoids with a wide range of sizes and relies on the supply of fresh tissue biopsy.^46-48^ The approach described in this research allows high-throughput production of AOAOs with defined size and consistent ciliation from *in vitro* expanded hABSCs, thus enables establishment of reproducible airway tissue models for effective diagnosis and therapeutic screening at a larger scale.

A recent study of enteroid engineered from intestinal epithelial stem cells suggests reversed epithelial polarity when transitioning the enteroid culture from Matrigel embedding to suspension.^49^ However, in our system, transferring the hABSC aggregates to Matrigel-embedded culture following as short as following one day in suspension culture remained exhibiting homogenous apical-out polarity. This indicates swift establishment of epithelial polarity in suspension culture of hABSC aggregates that remained stable following drastic change in their extracellular microenvironment.

The human lung is equipped with approximately 3 billion motile cilia, which are tiny hair-like protrusions (100-nm diameter) that constantly sweep mucus in the cephalic direction.^1^ Such nano-scale dimension combined with high beating frequency (10-20 Hz) makes measurement of cilia motility a challenging task requiring specialized high-speed video cameras at high magnification.^23,24^ As our work reveals, powered by cilia beating on its exterior surface, the AOAO exhibited persistent rotational motion when surrounded by Matrigel matrix. This novel phenomenon tackles the experimental and computational challenges associated with characterizing cilia motility and its pathophysiology, by enabling the conversion of nanoscale, high-frequency cilia beating into microscale, low-frequency AOAO rotation.

This study delivered a complete computational pipeline for quantifying organoid rotational motion from video data acquired using basic microscopic setup by leveraging the most recent developments in computer vision, specifically the tracking algorithms to conduct real-time analysis of video data. We have developed and deployed a computational framework that was able to extract significant features (such as rotational motion, percentage ciliation, and CBF) for characterization of airway organoids and their pathophysiology. The framework is generalizable and allows high-throughput feature extraction from video data, which enables processing of large quantities of data and robust statistical comparison. In this work, as a proof of concept for cilia motility analysis via AOAO rotation, we recorded and analyzed the angular velocity of the same AOAOs before and after the treatment with cilia beating inhibitors (EHNA and paclitaxel). The resulting significant deceleration of organoid rotation supported our hypothesis that the angular velocity of the AOAO correlates with and predicts the cilia motility.

Finally, our AOAO model recapitulates defective cilia motility under pathophysiological conditions and allows cilia injury assessment using patient-derived cells. We successfully engineering AOAOs from hABSCs derived from the PCD patient carrying *CCDC39* mutations and recapitulated the PCD-specific disease phenotype via analysis of AOAO rotational motion. This paves the way for using the AOAO rotation as a robust functional readout for developing effective assays for personalized disease management and therapeutic screening.

## Materials and Methods

### Materials

Detailed information regarding all reagents and equipment was described in Table 1, 2, 3, and 4.

**Table 1.** Reagents.

| **Name** | **Source** | **Catalog No.** |
| --- | --- | --- |
| A8301 (inhibitor of transforming growth factor $\beta$ kinase type 1 receptor) | Sigma-Aldrich | SML0788-5MG |
| BEBM™ Bronchial Epithelial Cell Growth Basal Medium | Lonza | CC-3171 |
| BEGM™ Bronchial Epithelial Cell Growth Medium<br>SingleQuots™ Supplements and Growth Factors | Lonza | CC-4175 |
| Bovine Collagen, Type I | Adbanced BioMatrix | 5225 |
| CHIR99021 (activator of WNT pathway) | Reprocell | 04000402 |
| Dulbecco's Phosphate-Buffered Saline (DPBS), 1X<br>without calcium and magnesium | Corning | 21-031-CV |
| DMH-1 (Inhibitor of BMP4/SMAD signaling) | Tocris Bioscience | 4126 |
| EHNA, hydrochloride (inhibitor of dynein) | Millipore Sigma | 32-463-010MG |
| Growth factor reduced Matrigel | Corning | CB 40230 |
| Heparin solution | Stemcell<br>Technologies | 07980 |
| HyClone™ FetalClone™ I Serum (a fetal bovine serum<br>alternative) | GE Healthcare | SH30080.03 |
| Hydrocortisone stock solution | Stemcell<br>Technologies | 07925 |
| Normal Human Bronchial Epithelial (NHBE) cells without<br>Retinoic Acid | Lonza | CC-2541 |
| Paclitaxel (inhibitor of microtubules) | Cayman Chemicals | 10461 |
| PneumaCult™-ALI Medium | Stemcell Technologies | 05001 |
| Penicillin-Streptomycin | Lonza Walkersville Inc. | 17-602E |
| RPMI 1640 with L-glutamine | Corning | 10-040-CV |
| Y27632 (Inhibitor of ROCKs) | Cayman Chemical | 129830-38-2 |
| 0.25% Trypsin-EDTA | Gibco™ | 25200056 |
| 96-well cell-repellent microplate with lid | GreinerBio-One | 655970 |

**Table 2.**
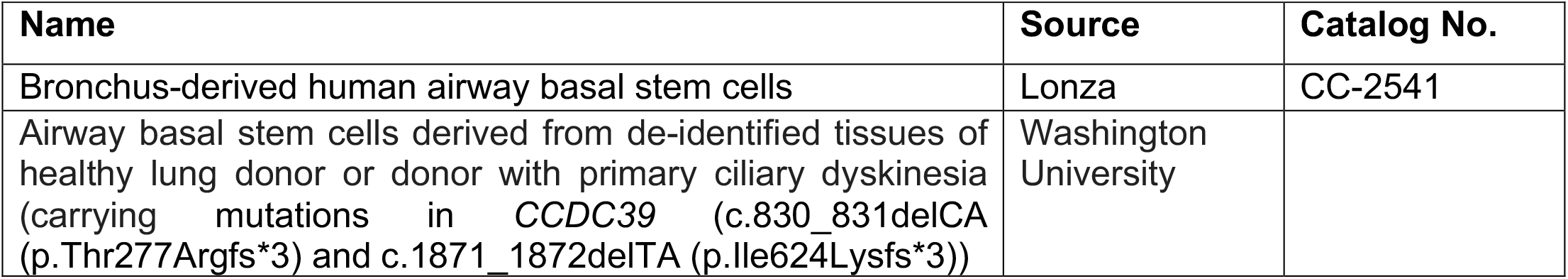
Cells.

**Table 3.**
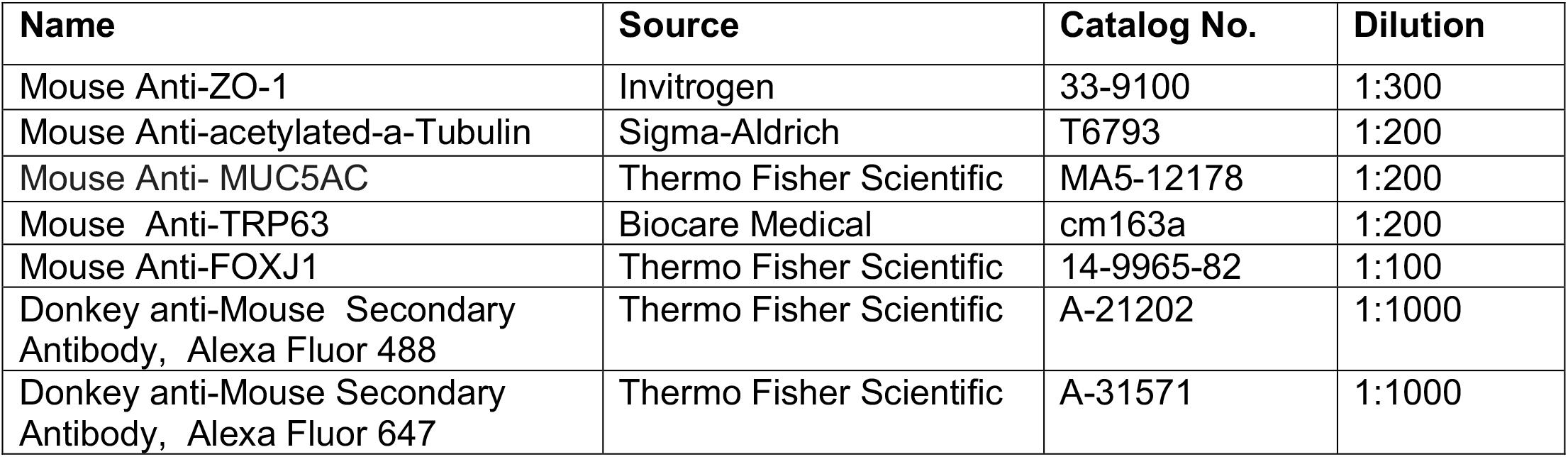
Antibodies.

**Table 4.** Equipment.

| Name | Source | Catalog No. |
| --- | --- | --- |
| EVOS FL Auto 2 Imaging System | Thermo Fisher Scientific | AMAFD2000 |
| Confocal Microscope | Zeiss | LSM 700 |
| Confocal Microscope | Zeiss | AxioObserver Z1 |
| High-speed sCMOS camera | PCO | pco.edge 5.5 |

### Methods

#### Culture of airway basal cells

Bronchus-derived human airway basal stem cells (hABSCs) were purchased from Lonza. Additional hABSCs were obtained from surgical excess of de-identified tissues of healthy lung donors or donors carrying mutations in *CCDC39* gene (c.830_831delCA (p.Thr277Argfs*3) and c.1871_1872delTA (p.Ile624Lysfs*3)) with permission of the institutional review board at Washington University in Saint Louis. The hABSCs were cultured in 804G-conditioned medium coated culture vessels in bronchial epithelial cell growth medium (BEGM) supplemented with 1 µM A8301, 5 µM Y27632, 0.2 µM of DMH-1, and 0.5 µM of CHIR99021 at 37°C with 5% CO_2_.^30^

#### Differentiation of airway basal cells into apical-out airway organoids

Human ABSCs (P2) were trypsinized and resuspended (5000 cells/ml) in differentiation medium (PneumaCult-ALI Medium) supplemented with 10 µM Y27632. 100 µL of resuspended hABSCs were placed per well in a 96-well cell-repellent microplate (GreinerBio-One, 655970). The cultures were maintained at 37°C with 5% CO_2_ for 21-28 days. To assess how IL-13 modulated organoid maturation, IL-13 (5 ng/mL) was supplemented to the differentiation medium during the entire differentiation period.

To assess organoid polarity in response to two-phase (ECM-deprived and then ECM-supported) culture, day-1 organoids formed in 96-well cell-repellent microplate were collected, resuspended in 40% (vol/vol) growth factor reduced (GFR) Matrigel and added to a new well plate that has been pre-coated with 40% GFR-Matrigel. The culture was then proceeded for an additional 20 days in differentiation medium.

#### Immunofluorescence Staining

Airway organoids were collected from the 96-well cell-repellent microplate and fixed with 4% paraformaldehyde (PFA) for 1 hour at 4°C. The fixed organoids were washed with PBS with 0.1% Tween-20 and permeabilized with 1% Triton-X for 1 hour before incubating with primary antibodies diluted in 1% bovine serum albumin (BSA) overnight. Next, the organoids were washed with PBS with 0.1% Tween20 and incubated with secondary antibodies. The nuclei were stained with 4′,6-diamidino-2-phenylindole (DAPI) before capturing z-stacks of stained organoids on a Zeiss LSM 700 laser scanning confocal microscope.

#### Scanning electron microscopy

The apical-out airway organoids (AOAOs) were fixed with 2.5% glutaraldehyde in 0.01 M PBS (pH 7.4) for 1 hour at room temperature. The organoids were washed 3 times in 0.01 M PBS and then post-fixed with aqueous 1% osmium tetroxide for 1 hour at 4°C. Next, the organoids were rinsed 3 times in 0.01 M PBS before dehydrating in a graded series of 30%, 50%, 70%, and 90% ethanol, followed by 3 changes in 100% ethanol. The organoids were further dehydrated in hexamethyldisilazane for 15 minutes and allowed to air dry. The fixed and dehydrated organoids were mounted on studs and sputter-coated with 5 nm gold-palladium alloy prior to imaging with JEOL JSM 7800.

#### Transmission electron microscopy

The AOAOs were rinsed in 0.01 M PBS and fixed with 2.5% glutaraldehyde in 0.01M PBS (pH 7.4) for 1 hour at room temperature. The organoids were washed 3 times in 0.01M PBS and then post-fixed with aqueous 1% osmium tetroxide containing 1% potassium ferricyanide for 1 hour at 4°C. Next, the organoids were rinsed 3 times in 0.01 M PBS before dehydrating in a graded series of 30%, 50%, 70%, and 90% ethanol, followed by 3 changes in 100% ethanol. The organoids were washed in Polybed 812 epoxy resin for 3 times for 1 hour each before polymerizing at 37° C overnight and then for additional 48 hours at 60° C. Finally, the prepared organoid samples were sectioned at 60 nm, placed on copper grids, and imaged with JEM 1400 Flash TEM.

#### Calculating percentage ciliation in AOAOs

Z-stack images of AOAOs, stained with Acetylated-α-Tubulin (Ac-α-Tub) and DAPI, were acquired using a Zeiss LSM 700 laser scanning confocal microscope. For percentage ciliation calculations, 3-4 cross-sections per organoids were selected from mid-z-stacks. The pixel coordinates of the edges of DAPI stained nuclei were used to determine the centroid of the apical-out organoid. By using k-means (k=1) clustering on the edge coordinates of DAPI-stained nuclei, we determined the centroid of the organoid in a robust and unsupervised manner.^50^ From the calculated centroid, each organoid cross-section was divided into 1-degree angular segments (Figure 1F). Finally, the presence of Ac-α-Tub immunofluorescence signal was detected in each angular segment. The angular segments containing the α-AcTub signal were recorded and used to calculate the percentage ciliation of the organoid.^51^

#### Calculating abundance of ciliated cells in AOAOs

Z-stack images of AOAOs, stained for the presence FOXJ1 using Anti-FOXJ1 antibody and for nuclei using DAPI, were acquired using a Zeiss LSM 700 laser scanning confocal microscope. For calculating the percentage abundance of FOXJ1^+^ ciliated cells, the mid-cross section of each organoid was selected from the z-stacks. The number of FOXJ1^+^ and DAPI^+^ cells were calculated using the Spots tool in the IMARIS software. The percentage of ciliated cells was calculated by normalizing the number of FOXJ1^+^ cells by the DAPI^+^ total cell number.

#### Matrigel embedding of AOAOs

Mature AOAOs at day-21 to day-28 of differentiation were collected together and embedded in Matrigel matrices. For Matrigel embedding, collected AOAOs were resuspended in 40% (vol/vol) GFR-Matrigel in differentiation medium, which was kept on a heat plate set to 37°C for 10 minutes to enable effective gelation. Upon matrix gelation, differentiation medium was added to the top of the AOAO-containing gel matrices. All matrix-embedded AOAOs were maintained at 37°C with 5% CO_2_. The next day, 30-seconds video recordings of AOAOs were captured using EVOS FL Auto 2 Imaging System.

#### Angular velocity calculation from video data

The video recordings of AOAOs were preprocessed by cropping to the region of interest containing the organoid, using Gaussian blur to reduce the noise, and smoothing variations in contrast to improve the performance of the tracking algorithm.^52,53^ Since the organoids generally have a spheroid shape, an ellipse was fit to the region of interest to mask out the surrounding region of the organoid. From the first frame of the video, a grid of equispaced correspondences was selected (Figure 3C). The correspondences were 5 pixels apart in each direction. These correspondences were tracked for the duration of the video by using the Lukas-Kanade (LK) tracking algorithm implementation from OpenCV python package.^32,33,54^ The distance covered by each correspondence was measured and converted to the rotational velocity.^55^ The correspondence which did not move, hence were marking the background, was filtered out. The rotational velocity was converted to angular velocity for each organoid by dividing by the distance of correspondence from the center (Supplementary Figure 4). To eliminate the error accumulation by the LK tracking algorithm over time, we recomputed the correspondences every 25 frames. The angular velocity for each organoid was the average angular velocity of all correspondences for the entire duration of the video. The difference in rotational and angular velocity was quantified by calculating the mean normalized deviation of velocity (rotational and angular) from the mean (Figure 3E) using the following equation:

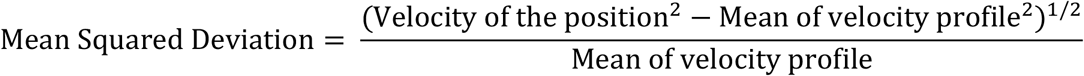

#### Treating AOAOs with Paclitaxel or EHNA for angular velocity analysis

Mature AOAOs (day-21 to day-28 of differentiation) were embedded in Matrigel for two days before treatment with desired chemical inhibitors of cilia motility. For paclitaxel, AOAOs were treated with paclitaxel (20 µM, diluted in differentiation medium) for 24 hours, with the control group being treated with an equal concentration of dimethyl sulfoxide (DMSO). For EHNA, AOAOs were treated with EHNA (0.1, 0.3, or 1 mM) for 2 hours, with the control group being treated with an equal concentration of phosphate buffer saline (PBS). Following chemical treatment of desire time periods, 30-seconds video recordings of AOAOs, pre- and post-treatment, were captured using EVOS FL Auto 2 Imaging System.

#### Imaging the AOAO cilia beating for kymographs generation and cilia beating frequency (CBF) calculation

Mature AOAOs (day-21 to day-28 of differentiation) were transferred to a 1.5-mL Eppendorf tube and kept on ice. Cold collagen type 1 was neutralized with the neutralization solution, added to the organoids at a final concentration of 2 mg/mL, and the entire AOAO-collagen mixture was transferred to glass bottom region of a Mattek dish. The Mattek dish containing organoids in collagen was kept on ice for additional 10 minutes until the organoids settle down to the bottom of the dish. The Mattek dish was then kept on a heat plate set to 37°C for 10 minutes. 1 mL of differentiation medium was placed in the dish before capturing video recording of cilia beating using on a Zeiss AxioObserver Z1 microscope with a 100X, 1.45 NA objective and pco.edge 5.5 camera. The high-speed video recordings of cilia beating were preprocessed using the previously described method to smoothen noise and variations in contrast. Additionally, the Contrast Limited Adaptive Histogram Equalization (CLAHE) function in Python v. 3.7 software was used to improve the contrast of the cilia with respect to the background.^53,56^ From preprocessed video, the region of interest on organoid surface containing cilia was cropped. The normal vector with respect to the organoid surface for each region of interest was calculated. The pixel intensity along the normal vector was then mean pooled for each frame, thereby generating the kymograph for ciliary motion. The peaks in the kymograph were counted and divided by the duration of the video to obtain the CBF value for the organoid. At least 10 kymographs were generated per organoid and the average value represented the CBF of the organoid.

### Statistics

Quantitative data were displayed as means ± s.d. Statistical significances were determined using one-way analysis of variance (ANOVA) with post-hoc Tukey’s Test and unpaired t-test. Statistical analyses were performed using GraphPad.

### Graphics

Schematics were created using Microsoft PowerPoint and Biorender. Plots were prepared using GraphPad.

## Author Contribution

† These authors contributed equally. P.W., P.Y., A.B.F. and X.R. contributed to the overall design of the project, data interpretation, and preparation of the manuscript. P.W. and Y.L. performed the experiments and analyzed data. L.A.P. and S.C.W. performed image acquisition regarding cilia beating. D.B.S and J.M.F. performed electron microscopy studies. E.R.J. assisted with figure preparation. S.L.B., A.H. and J.X. provided patient-derived human airway basal stem cells and advised on experiments related to these cells.

## Acknowledgement

We are grateful to Misti West for laboratory management, and to Ming Sun and Mara L. Sullivan from the Center for Biologic Imaging at University of Pittsburgh for assistance with TEM sample processing.

## List of Supplementary Information

**Supplementary Figure 1.**
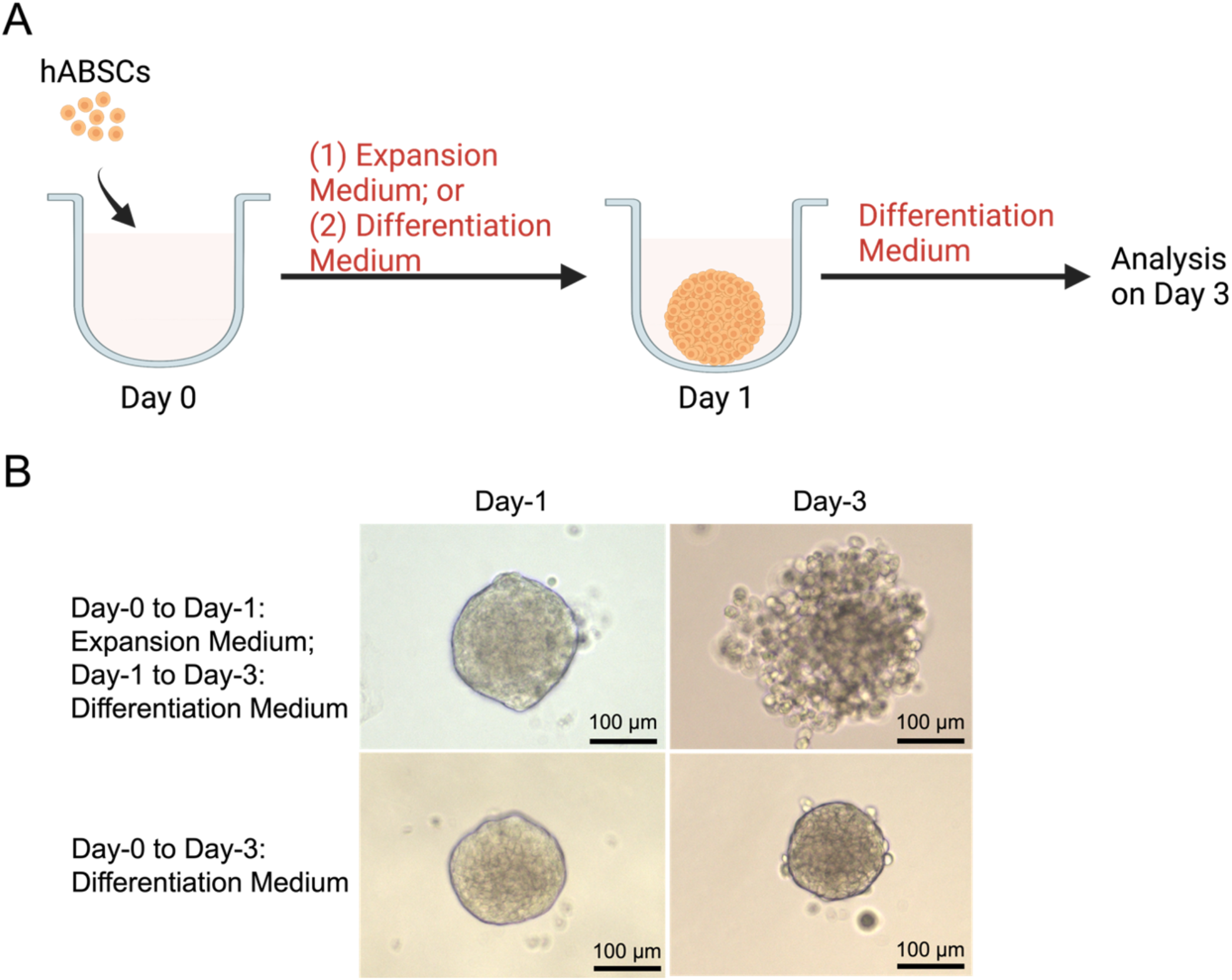
Airway organoid formation in different medium in 3D suspension culture. (A) Diagram showing the conditions under comparison: during day-0 to day-1 of 3D suspension culture, the organoid was either cultured in BEGM-based expansion medium or in differentiation medium (PneumaCult-ALI); and during day-1 to day-3 of culture, the organoid was cultured in differentiation medium. (B) Brightfield images of organoid integrity on day-1 and day-3 of culture.

**Supplementary Figure 2.**
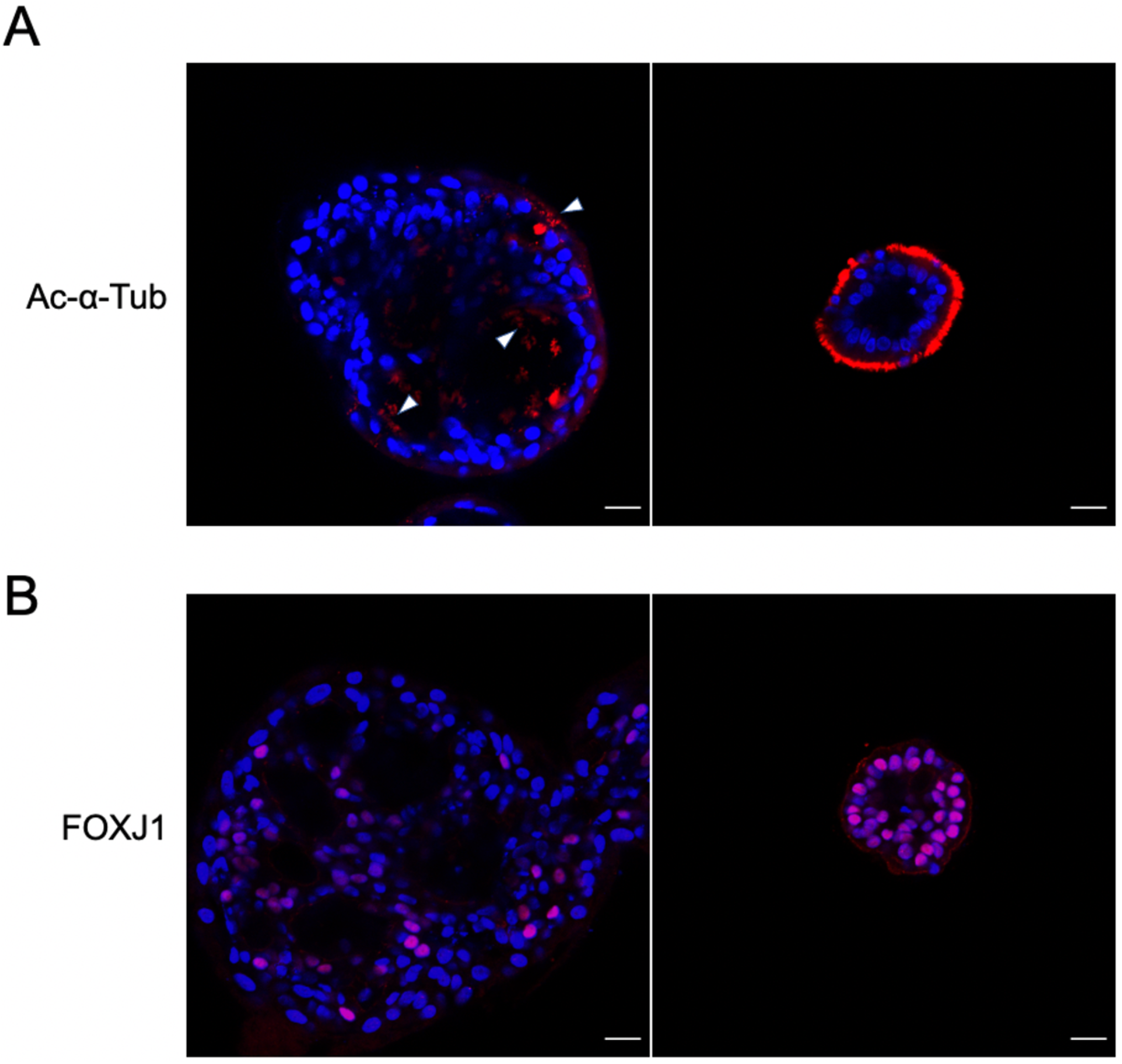
Large organoid bodies in the organoid culture transferred to Matrigel-embedding following 1 day in suspension from dissociated hABSCs. (A,B) Immunofluorescence staining of Ac-α-Tub (A) and FOXJ1 (B) in both large organoid bodies and regular-sized organoids on day-21 of culture.

**Supplementary Figure 3.**
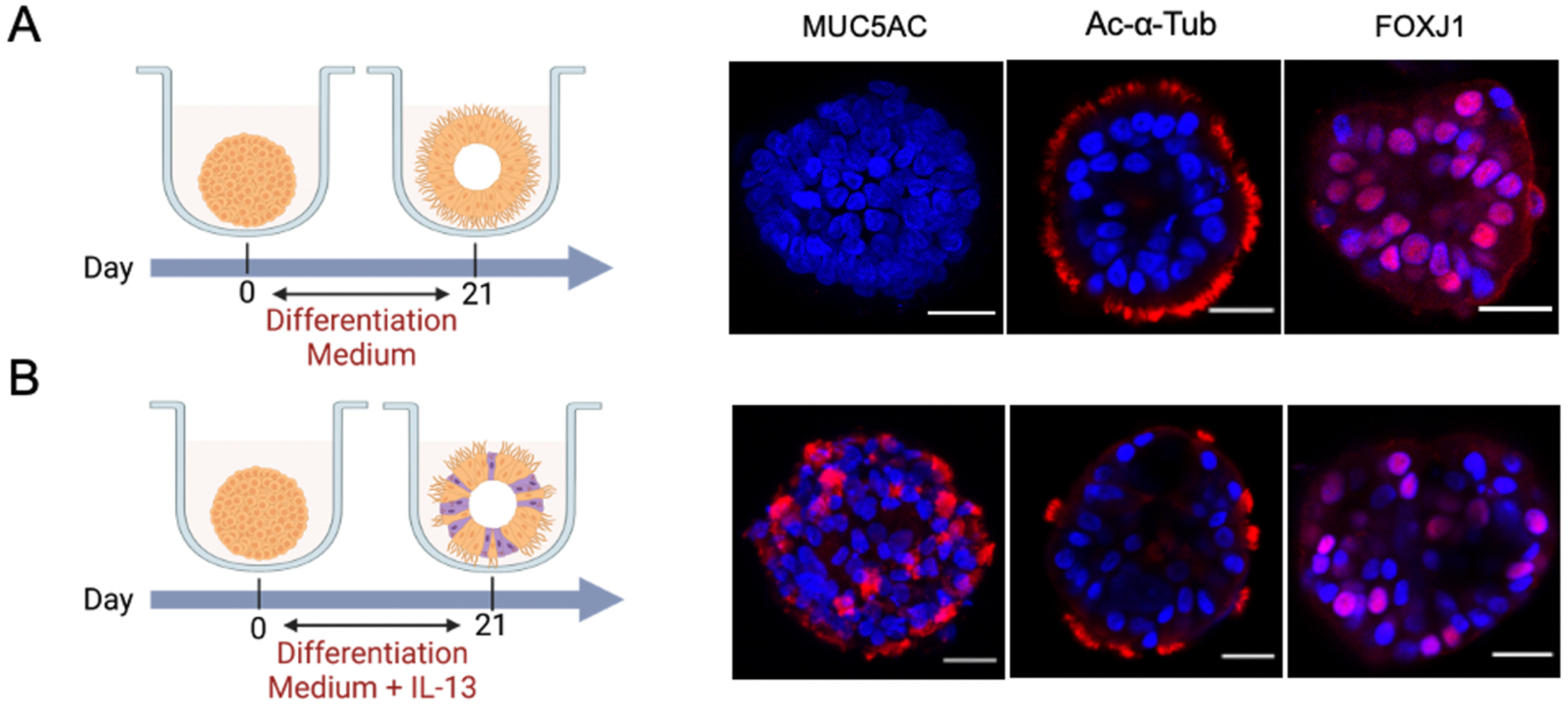
Characterizing cellular composition in AOAOs differentiated in the presence and absence of IL-13. (A,B) Immunofluorescence staining of MUC5AC, Ac-α-Tub and FOXJ1 in AOAOs following 21 days culture in either differentiation medium (A) or in differentiation medium supplemented with IL-13 (5 ng/mL).

**Supplementary Figure 4.**
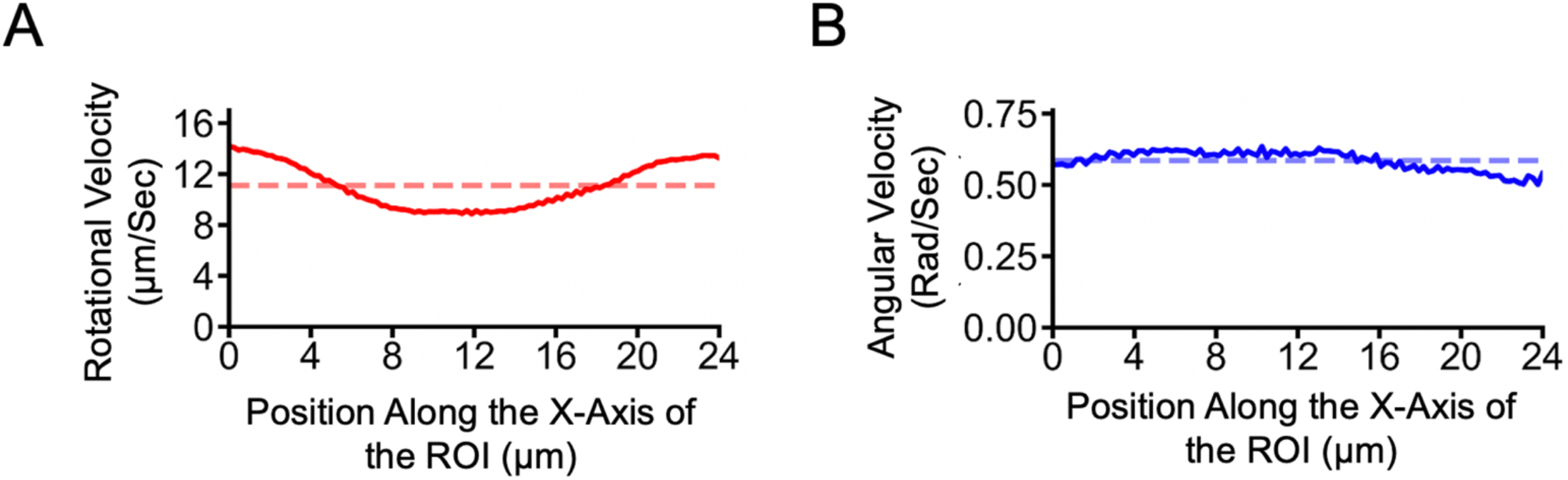
Rotational (A) and angular (B) velocity profiles of AOAO rotation. The solid trace represents the average velocity of all correspondences on the organoid perpendicular to that position on the X-axis of the ROI. The dashed line represents the mean velocity profile.

**Supplementary Video 1**. Day-21 AOAO from 3D suspension culture with cilia beating on its exterior surface.

**Supplementary Video 2**. Matrigel-supported rotation of mature AOAO. Mature AOAO formed in 3D suspension culture was transferred to 40% GFR-Matrigel embedding, where the exterior-facing cilia propelled against the Matrigel matrix and drove the organoid to rotate.

**Supplementary Video 3**. Demonstration of computer vision framework for tracking the rotational motion of AOAOs embedded in Matrigel. The video showed the tracking of organoid rotation for 5 seconds comprising the 4 steps as described in Figure 3D. The first panel showed the raw organoid video with extracted ROI and the fit ellipse to suppress background noise. The second panel showed the implementation of the tracking algorithm and visualization of the trajectory of each correspondence as green trace. The third panel was time-synchronized plot of the instantaneous angular velocity of the organoid. The correspondences in panel 2 were recomputed every 1 second to minimize the error accumulation by tracking algorithm. The traces for the trajectory of each correspondence were reset after 1 second and correspondences were recomputed from the new orientation of organoid. The rotational velocity for each of the 1 second segments of the organoid video was averaged to find the rotational velocity of the organoid over the duration of entire video.

**Supplementary Video 4**. Visualization of EHNA modulation of AOAO rotation. The first row showed the rotational motion of mature AOAO before EHNA treatment, and the second row showed the same organoid following 2 hours of EHNA treatment. The first column showed treatment with 0 mM of EHNA (control), and the second column showed treatment with 1 mM of EHNA.

**Supplementary Video 5**. Tracking the rotational motion of mature AOAOs engineered from healthy and PCD (CCDC39 mutation) hABSCs. The first row showed the healthy AOAO, its tracking demonstration, and the corresponding time-synchronized angular velocity plot. The second row showed the PCD AOAO, its tracking demonstration, and the corresponding time-synchronized angular velocity plot.

